## Supplementary Material for "Semantic novelty modulates neural responses to visual change across the human brain"

**This File includes:**

Figs. S1 to S24
Tables S1 to S12
Equation S1
References (1 to 12)

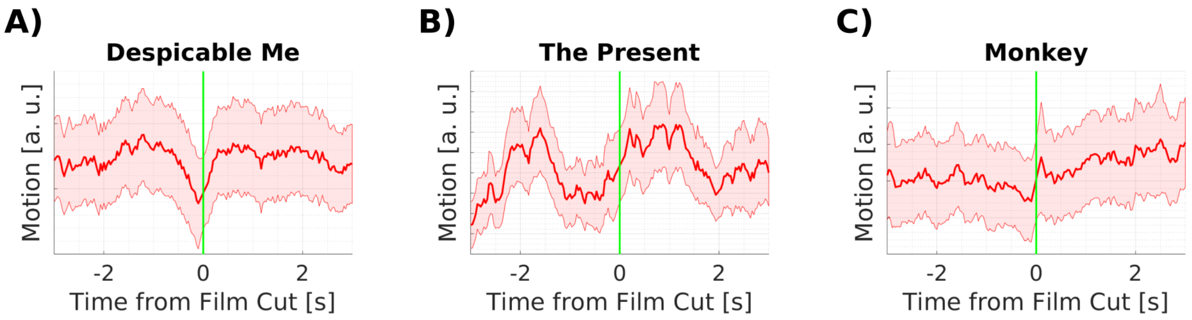

**Figure S1: Motion around film cuts differs for each video.** A) Motion in ‘Despicable Me’ decreases before film cuts, likely driving the effect in Figure 1F. Therefore, it is likely that professional editors decrease motion before cuts. B) Motion in ‘The Present’ peaks about 2 seconds before and 1.5 seconds after film cuts. Two film cuts causing an artifact have been removed for this figure. The artifact was caused by a large peak of motion between these cuts. C) Motion around film cuts in the monkey movies, does not show as much change as Despicable Me and The Present.

| Patient ID | Age | Sex | Notes |
|---|---|---|---|
| Pat_01 | 58 | M | prior laser ablation in right hemisphere mesial temporal lobe |
| Pat_02 | 22 | M | prior resection on right frontal lobe |
| Pat_03 | 33 | F | |
| Pat_04 | 46 | F | |
| Pat_05 | 48 | M | |
| Pat_06 | 36 | F | |
| Pat_07 | 43 | M | Spanish native, good English language comprehension |
| Pat_08 | 41 | F | |
| Pat_09/Pat_09_02 | 51 | M | |
| Pat_10 | 24 | M | Spanish native |
| Pat_11/Pat_11_02 | 37 | M | Spanish native, no English language comprehension |
| Pat_12 | 52 | F | |
| Pat_13 | 24 | M | |
| Pat_14 | 20 | M | |
| Pat_15 | 56 | F | undergone stereo EEG for neuropathic pain |
| Pat_16 | 43 | F | |
| Pat_17 | 27 | F | Spanish native, low English language comprehension |
| Pat_18 | 28 | F | Lesion in Occipital cortex with interictal activity; vision in right eye affected |
| Pat_19 | 46 | M | |
| Pat_20 | 35 | F | |
| Pat_21 | 48 | M | |
| Pat_22 | 19 | M | |
| Pat_23/Pat_23_03 | 36 | F | |

**Table S1: Patient demographics.** Data from patients 09, 11 and 23 comes from reimplants at different times.

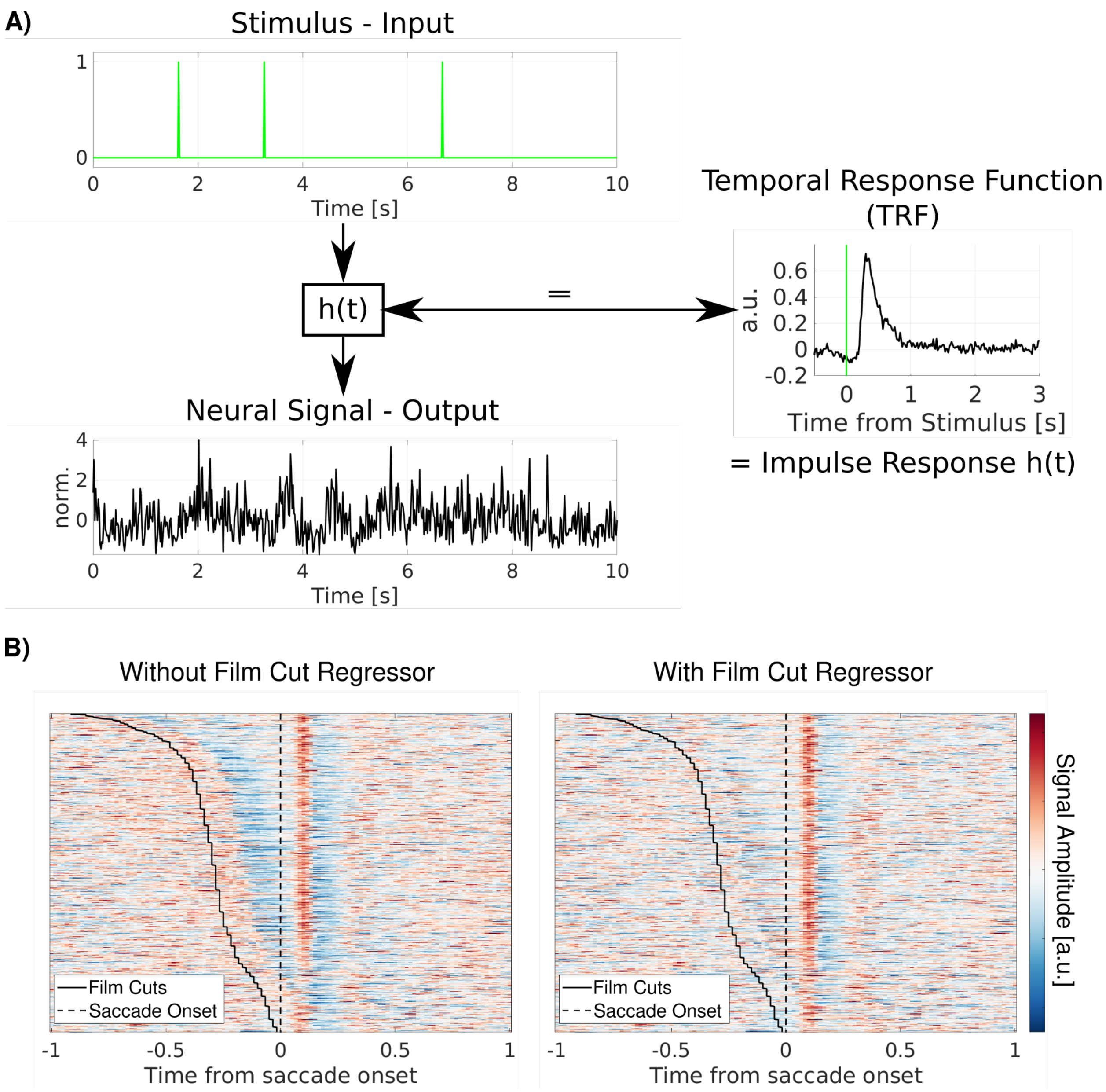

**Figure S2: Neural responses are identified through a linear system identification approach. A)** Motion, saccade or film cut onset are considered inputs to a linear system. The neural signal is considered the output. The temporal response function is the filter that maps from the input to the output through a convolution B) Including multiple regressors as inputs to the system allows to control for linear interactions. Left: Saccade evoked responses in the raw LFP in a channel in the occipital lobe. Each line corresponds to the neural signal within a time window of 2s around a single saccade. Film cuts occur within a second before saccades and would confound the average. Right: Including scene cuts in the model removes their linear contribution to the saccade evoked responses[1,2].

**Equation S1: Ridge regression.** The temporal response functions $h$ describes the linear mapping from input signal $S$ to the neural response $r$ :

$$r = Sh + n,$$

where $h$ is a vector of size T, $S$ is the input signal (regressor) arranged as a toeplitz matrix of size *N* by *T*, with *N* time samples by *T* delays, $r$ is the observed neural signal as a vector of length *N*, and $n$ is the residual unexplained noise. This temporal response function can be estimated through ordinary least squares, which we regularize using ridge regression:

$$h = \left(S^T S + \gamma \lambda\right)^{-1} S^T r \ .$$

Here $\lambda$ is the mean eigenvalue of the covariance matrix $S^T S$. We chose regularization constant $\gamma = 0.3$.

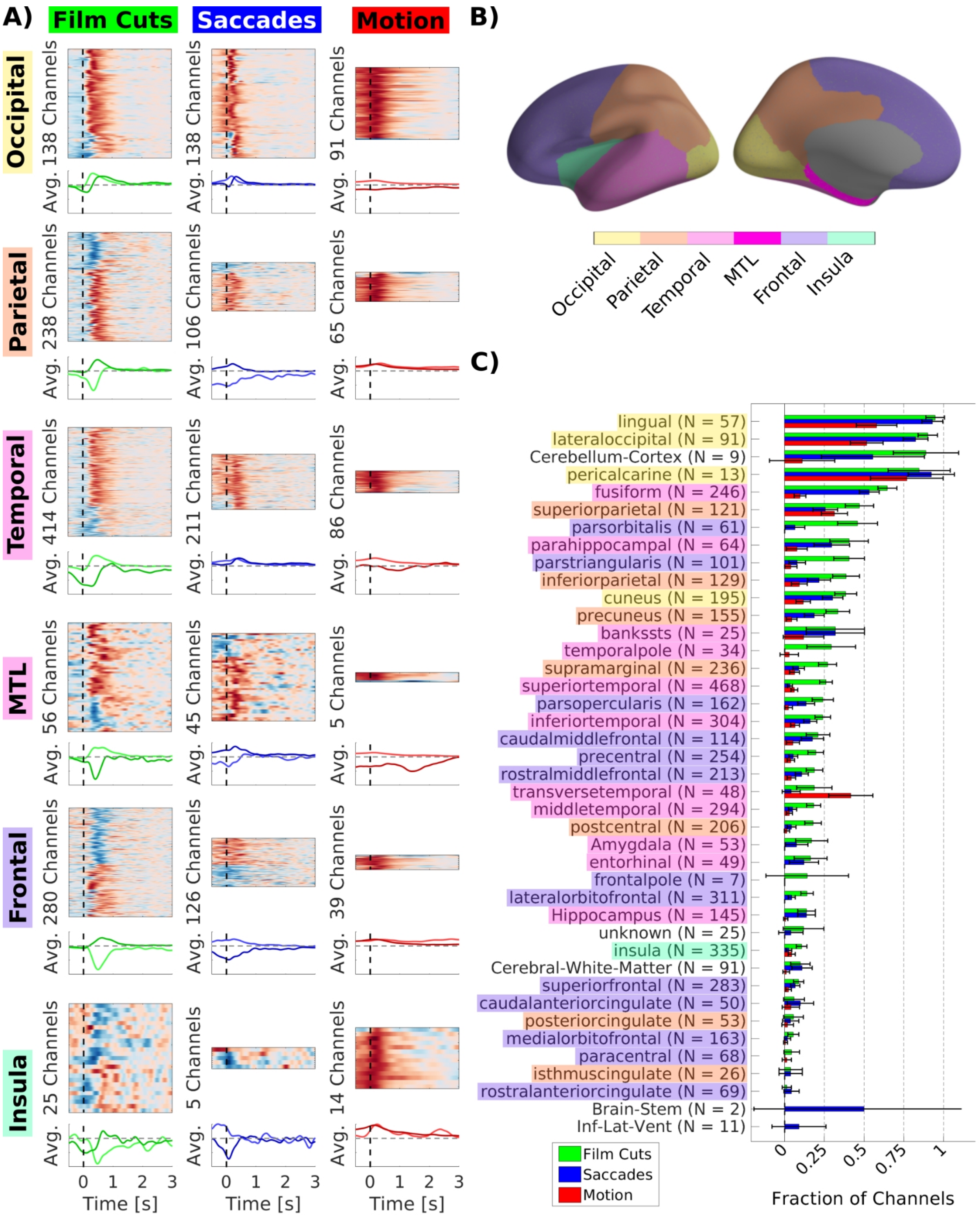

**Figure S3: Most channels respond to film cuts and saccades and only few to motion.** A) TRFs for channels with significant responses to film cuts, saccades and motion in all brain areas. TRFs are smoothed with a Gaussian window with a standard deviation of 53ms. B) Definition of lobes in A. Case courtesy of Assoc Prof Frank Gaillard, Radiopaedia.org, rID: 46846 C) Ratio of responsive channels out of total channels in each area. Brain areas are localized by the Desikan-Killiany and Aseg atlas in freesurfer[3,4]. When only one channel is in a given anatomical area the channel pair is counted as 0.5 channels in the given area. Error bars depict the 95% confidence interval of the proportion of responsive channels. As a caveat, note that this finer parcellation assumes good anatomical alignment with the atlas, which may not always be the case.

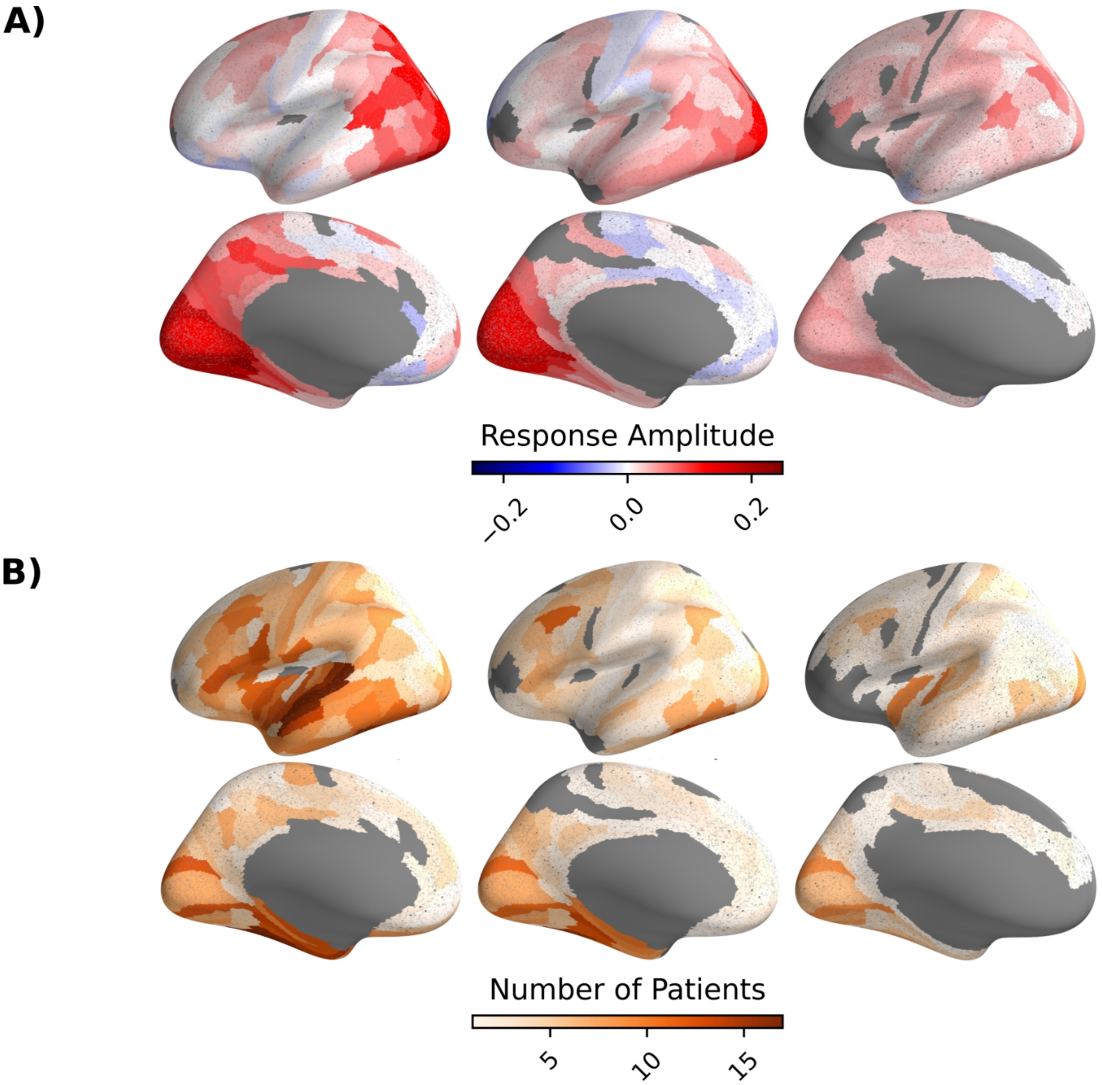

**Figure S4: Responses to film cuts and saccades are more consistent across patients than responses to motion.** A) Weighted average of response amplitude as in Figure 2B. B) Number of patients with significant electrodes in each parcel.

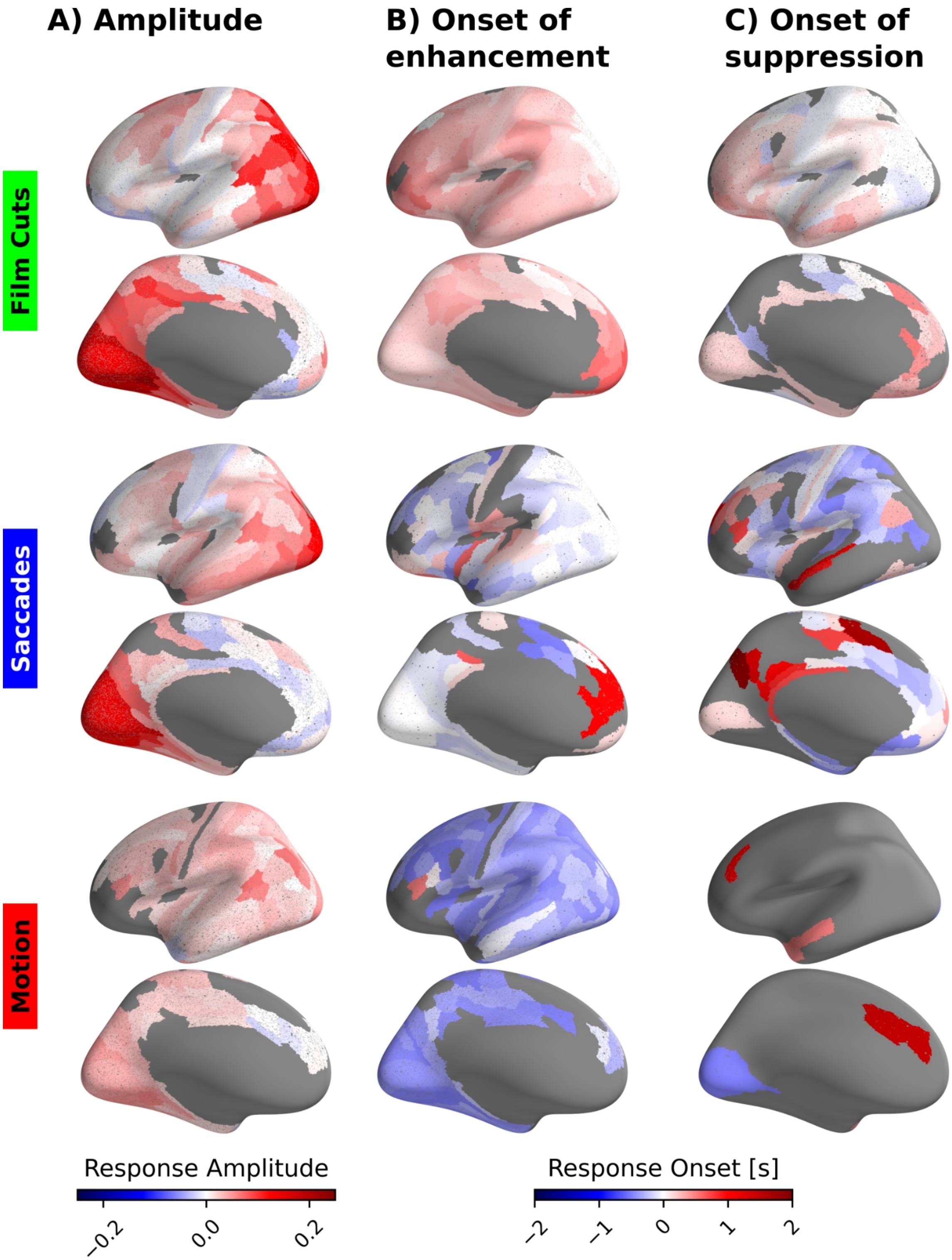

**Figure S5: Film cuts and saccades exhibit specific profiles of the onset of responses.** A) Weighted average amplitude of responsive channels in each parcel of the Glasser atlas. Red colors indicate an enhancement of the amplitude of BHA over baseline, blue colors suppression over baseline. Amplitudes are calculated at the extrema (maximum or minimum of the response). B) Weighted average onset of responses. Only channels with enhancement of BHA are considered. Red colors indicate a delay after the stimulus, i.e. the response starts after the stimulus. Blue colors indicate responses starting before the stimulus onset. Responses to film cuts start predominantly after the cuts, with increasing delays toward frontal areas. Responses to saccades start close to saccade onset in visual areas and before saccade onset in most other brain areas. Responses to motion are predictive of motion in most channels. Onset has been estimated as the full-width-half-max of the TRF around the peak. C) Only channels with suppression of BHA are considered. Responses in parts of the occipital and superior temporal lobe are predicitve of cuts, while the onset of responses in the frontal lobe lags cuts. Suppression of BHA in the case of saccades is mostly predictive of saccades. While many channels show predictive responses, they are partly explained by smoothing of the temporal response functions through regularization.

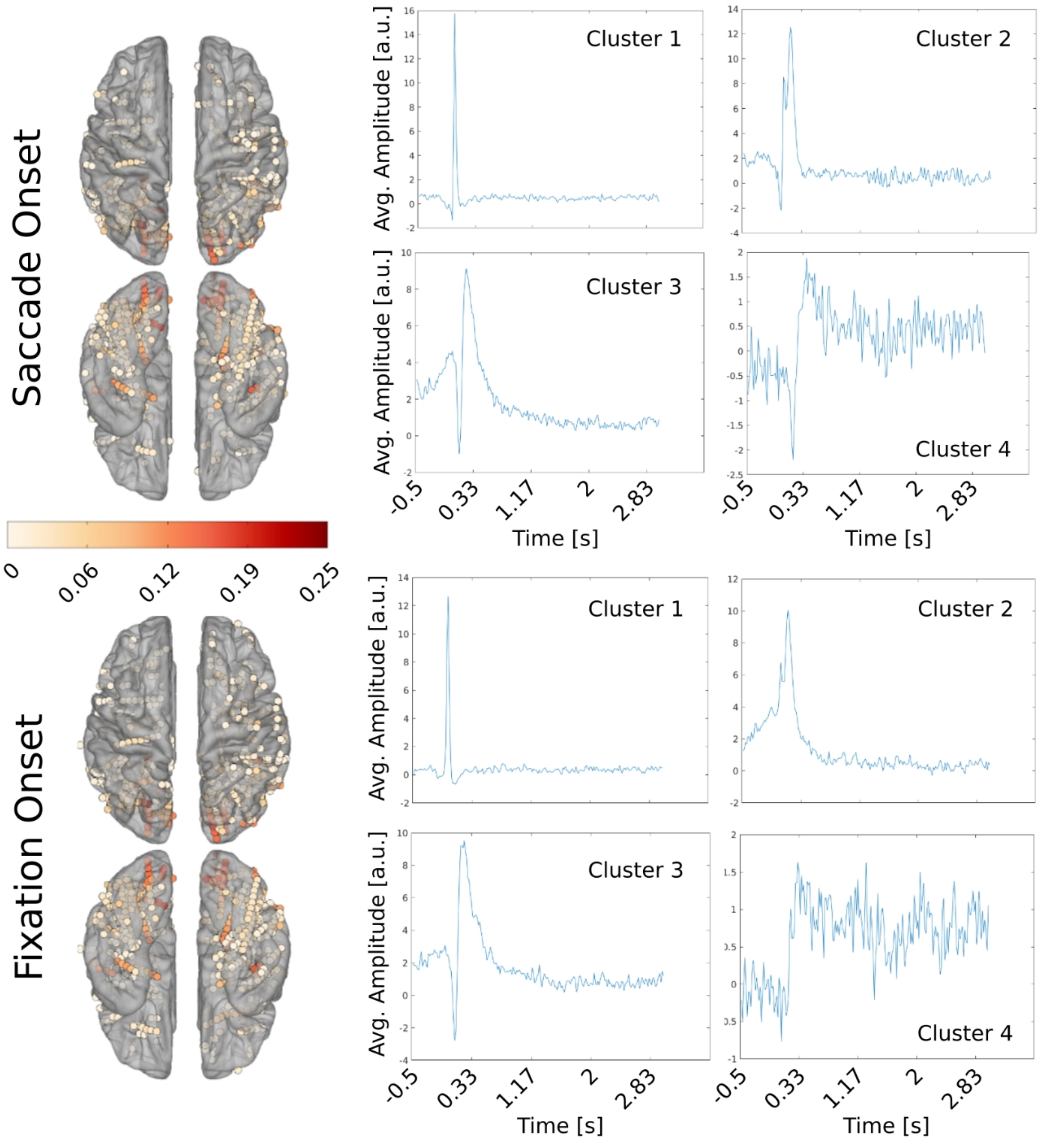

**Figure S6: Responsive electrodes and average filters are similar if analysis is conducted for saccade or fixation onset.** Temporal response functions for each channel were identified as in Figure S2. Then the stimulus vector was convolved with the temporal response function to predict the signal in each channel. The color of electrodes in the spatial plots on the left indicate the correlation of the predicted signal with the original signal in each significant channel. Significance was measured by permutation statistics similar to Figure 2. A surrogate distribution of filters was identified by shuffling the timing of the neural signal in relation to the stimulus vector. The correlation value obtained by correlating the original filter was then tested against the correlation values obtained from the shuffled filters. We visualize channel locations on the freesurfer fsaverage brain[5] with the iELVis MATLAB toolbox[6]. Panels on the right side of the figure show average temporal response functions in clusters of correlated responses[7]. The same analysis was conducted for a stimulus vector with impulses at saccade onset (top) and fixation onset (bottom). Both the spatial extent of significant electrodes and the temporal response functions are similar in both analyses.

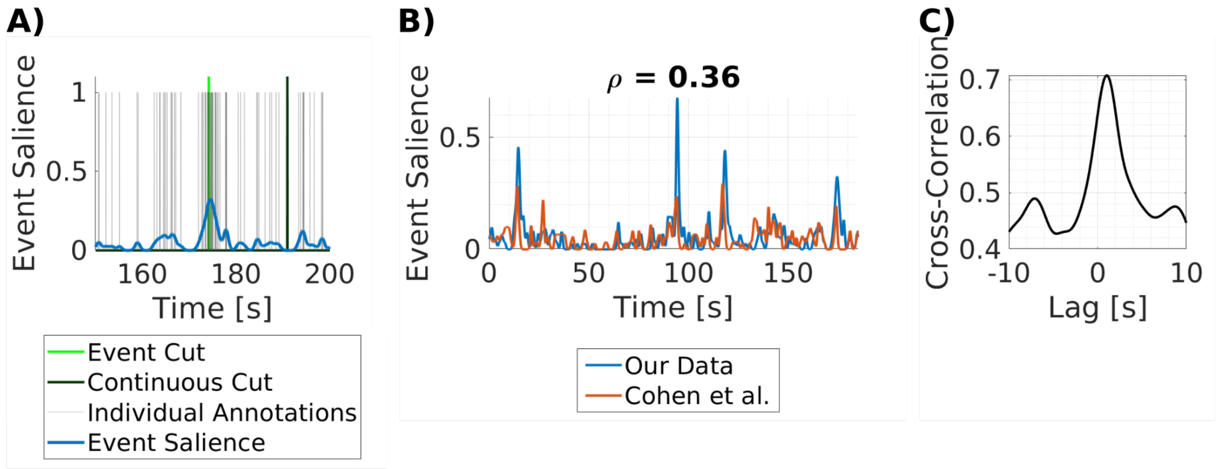

**Figure S7: Event boundary annotations correspond with film cuts and are reliable across datasets.** Event boundary annotations have been collected from a population of 63 participants recruited on prolific.com. Participants watched one of the movies from the dataset and responded with a key press when they felt that there was a change in the narrative. A) Light gray lines indicate the timing of individual event boundary responses across all participants. The regressor of impulses at the time of responses is smoothed with a Gaussian kernel with a standard deviation of 0.5s (blue line). The resulting regressor is a measure of the salience of the event. The event salience corresponding to film cuts can be determined by this event salience regressor. For each video event film cuts are defined as the film cuts with the highest event salience, at the peaks of the event salience regressor (bright green line). Continuous cuts are cuts at times of low event salience and are matched to event cuts in their low level visual features (dark green line). B) Event salience from our data is similar to event salience measured by Cohen et al.[8]. Cohen et al. recorded event boundary annotations from 21 participants. Event boundary annotations were smoothed with a Gaussian kernel with a standard deviation of 0.5s. C) Cross-correlation of event salience from our data and from Cohen et al. peaks at a delay of ~0.66s.

| Video | N | excluded | Total Cuts | Event Cuts |
|---|---|---|---|---|
| Despicable Me English | 73 | 10 | 157 | 9 |
| Despicable Me Hungarian | 20 | 0 | 160 | 11 |
| The Present | 26 | 2 | 56 | 3 |
| Monkey 1 | 29 | 3 | 54 | 8 |
| Monkey 2 | 27 | 3 | 63 | 15 |
| Monkey 5 | 25 | 2 | 71 | 11 |
| **Total** | **200** | **20** | **561** | **57** |

**Table S2: Event annotation data.** 'N': total number of participants recruited online to annotate event boundaries. 'excluded': number of participants excluded from analysis because either no boundaries were annotated or the attention test failed. The attention test consists of white boxes flashing on a black background. 'Total Cuts': Total number of film cuts in each video. 'Event Cuts': number of event boundaries and matched continuous cuts selected.

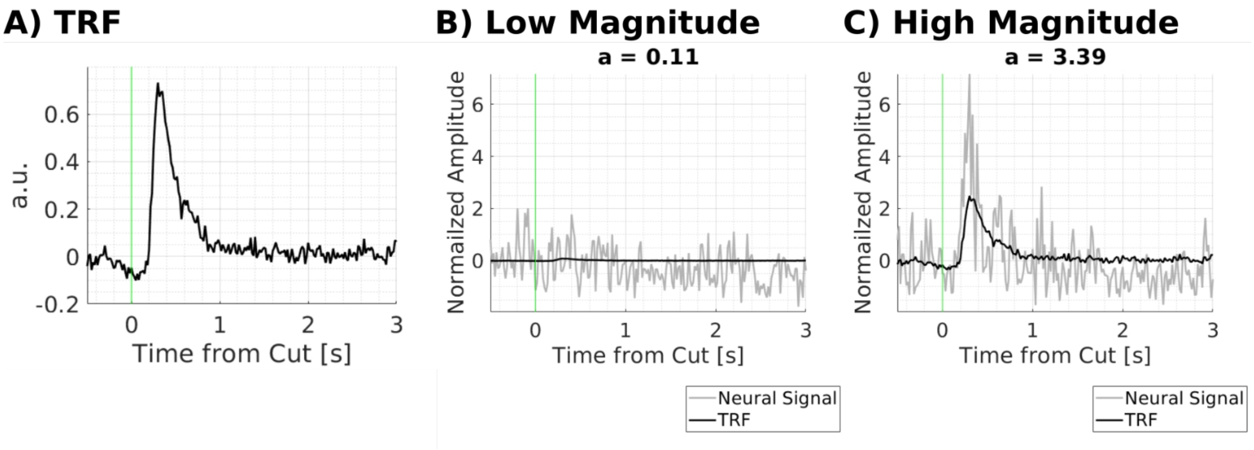

**Figure S8: Estimation of the magnitude of responses to individual film cuts and saccades.** A) Temporal response functions are estimated on all stimuli in each channel. TRFs are then fit to the neural signal around each event in the stimulus in the same time window. The magnitude is estimated as the coefficient ‘a’ that the TRF is multiplied with to best fit the neural data. B) For low magnitude responses a < 1. C) For high magnitude responses a > 1.

| **Factor** | **d.f.** | **F** | **p-value** |
|---|---|---|---|
| Condition | 1 | 33.83 | **$6.9*10^{-9}$** |
| Region | 5 | 5.75 | **$2.8*10^{-5}$** |
| Patient | 25 | 5.23 | **$9.2*10^{-16}$** |
| Error | 2270 | | |

**Table S3: Results of mixed-design ANOVA testing the significance of differences in response amplitudes to event versus continuous cuts (Figure 3A).** Condition is the fixed effect of amplitudes of event or continuous film cuts. Region is the fixed effect of brain region. Patient is a random effect.

| Factor | d.f. | F | p-value | | Factor | d.f. | F | p-value |
|---|---|---|---|---|---|---|---|---|
| **Occipital lobe** | | | | | **MTL** | | | |
| Condition | 1 | 0.84 | 0.36 | | Condition | 1 | 23.24 | **$5.4*10^{-6}$** |
| Patient | 12 | 2.18 | **0.01** | | Patient | 15 | 1.28 | 0.23 |
| Error | 262 | | | | Error | 95 | | |
| **Parietal lobe** | | | | | **Frontal lobe** | | | |
| Condition | 1 | 0.23 | 0.64 | | Condition | 1 | 1.05 | 0.30 |
| Patient | 19 | 4.93 | **$1.3*10^{-10}$** | | Patient | 23 | 1.96 | **0.005** |
| Error | 455 | | | | Error | 535 | | |
| **Temporal lobe** | | | | | **Insula** | | | |
| Condition | 1 | 48.45 | **$7*10^{-12}$** | | Condition | 1 | 0.66 | 0.42 |
| Patient | 23 | 3.65 | **$2.4*10^{-8}$** | | Patient | 11 | 3.24 | **0.003** |
| Error | 803 | | | | Error | 37 | | |

**Table S4: Results of mixed-design ANOVA testing the significance of differences in response amplitudes to event versus continuous cuts within each brain region (Figure 3A).** Condition is the fixed effect of amplitudes of event or continuous film cuts. Patient is a random effect.

| | Occipital | Parietal | Temporal | MTL | Frontal |
|---|---|---|---|---|---|
| **Median Jaccard distance** | 0.31 | 0.71 | 1 | 1 | 1 |
| **p-value** | 0.999 | 0.4 | **0.005** | **0.032** | **0.025** |
| **N** | 12 | 9 | 22 | 11 | 14 |

**Table S5: Results of permutation statistics to test specificity of responses to event and continuous film cuts (Figure 3D).** 1,000 permutations. FDR corrected at q = 0.05. N: number of patients

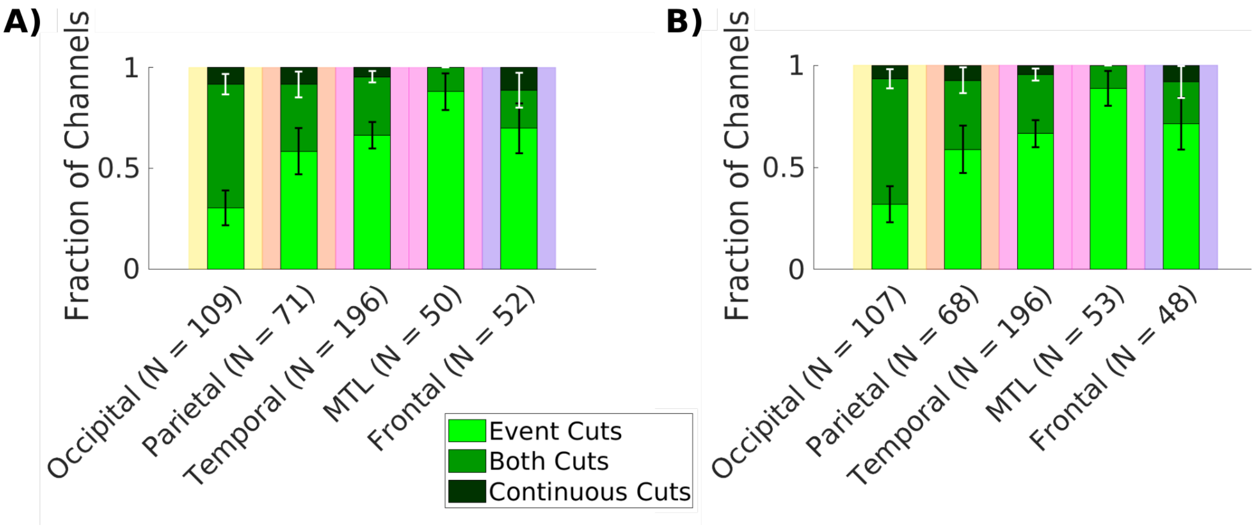

**Figure S9: TRFs for event cuts and continuous cuts do not depend on motion.** A) TRF model includes event cuts, continuous cuts and saccades. Fraction of channels out of responsive channels with significant responses to event cuts only (bright green), continuous cuts only (dark green), or both event and continuous cuts (mixed green). B) Same as panel A with a TRF model including event cuts, continuous cuts, saccades and motion.

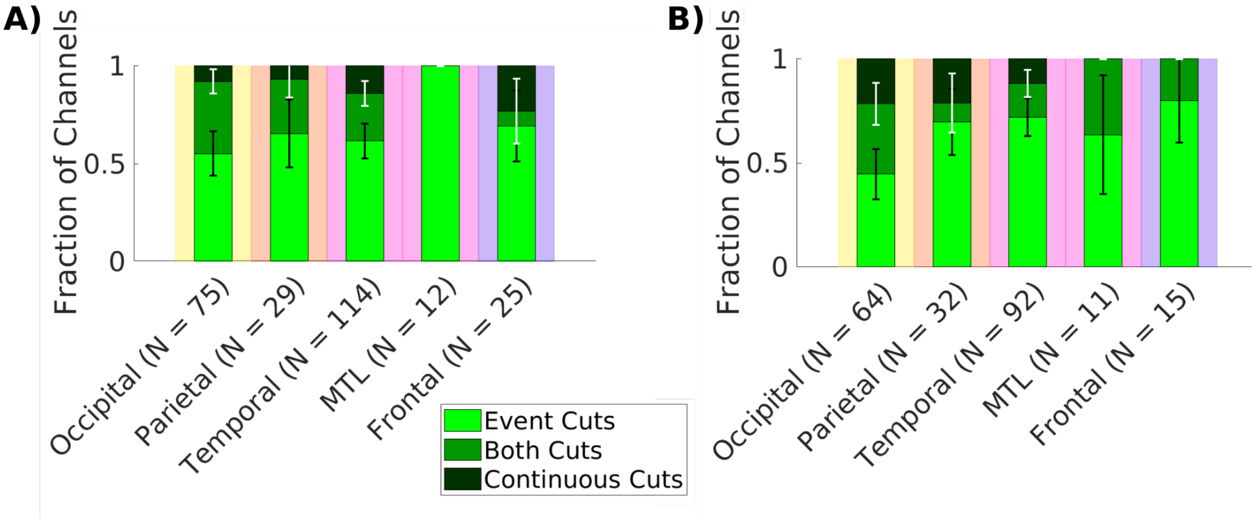

**Figure S10: Event cuts dominate neural responses for comics and monkey videos.** Analysis of event cuts and continuous cuts for Monkey videos (‘Monkey1’, ‘Monkey2’, ‘Monkey5’, 15min total) and comics (‘Despicable Me English’, ‘Despicable Me Hungarian’, ‘The Present’, 28.6min total). A) Monkey videos only. Fraction of channels out of significant channels responding to event cuts only (bright green), continuous cuts only (dark green), or both event and continuous cuts (mixed green). B) Same analysis as panel A, but for data of comics.

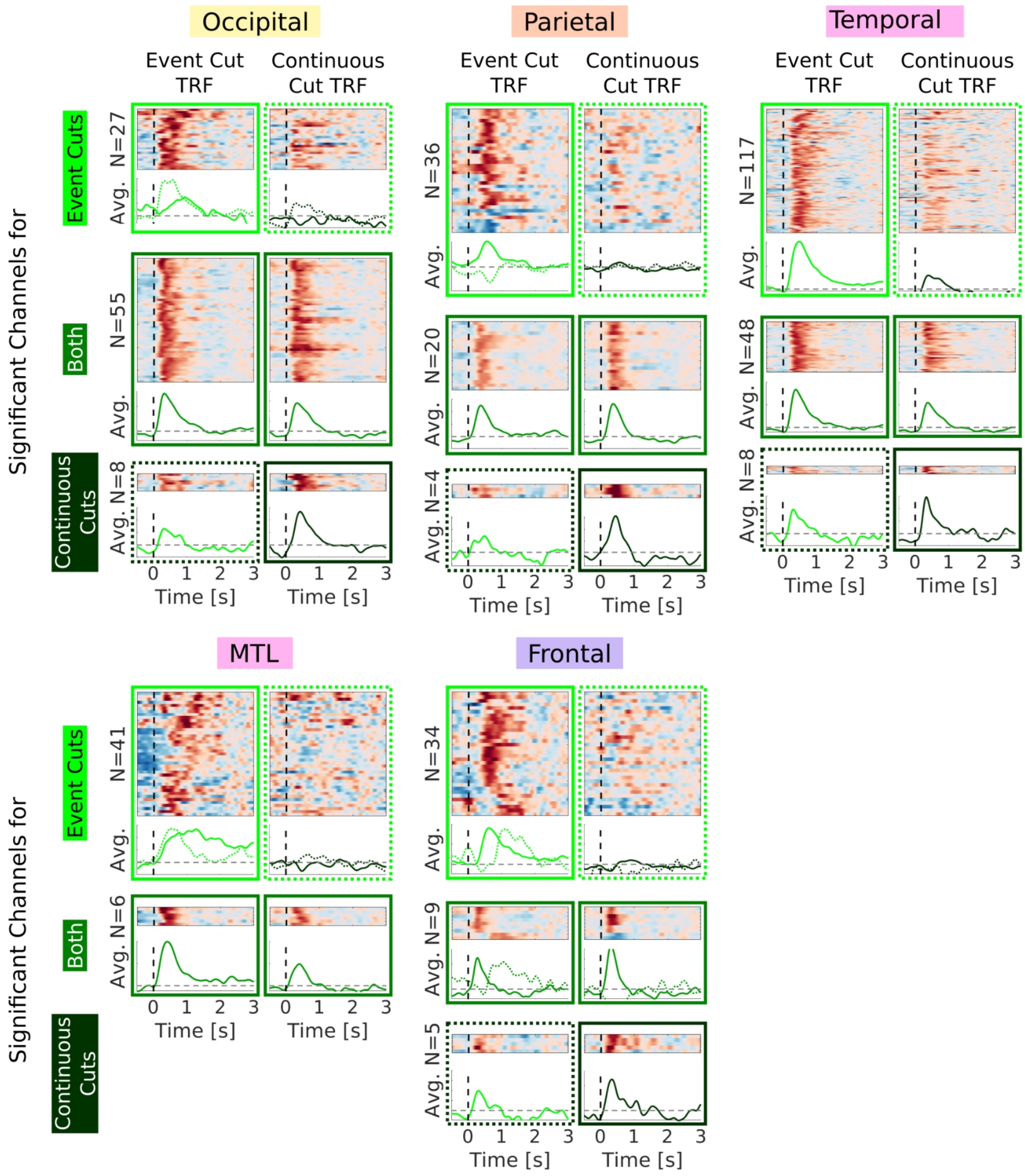

**Figure S11: More channels respond to event cuts than continuous cuts.** Separate filters are determined for event cuts and continuous cuts (Figure 3C). In the top of each box normalized temporal response functions are depicted. The bottom of each box shows average temporal response functions. Averages are computed for clusters of similar temporal response functions. Bright green boxes on top show channels with significant responses to event cuts only. Boxes with solid lines indicate the significant TRFs to high event cuts in these channels. Boxes with dashed lines indicate non-significant TRFs to continuous cuts in the same channels. Dark green boxes on the bottom show TRFs in channels with significant responses to continuous cuts. Boxes with solid lines, here, show the significant TRFs to continuous cuts in these channels. Boxes with dashed lines show the non-significant TRFs to event cuts in the same channels. Mixed green boxes in the middle show channels with significant TRFs to both event and continuous cuts. TRFs are grouped in lobes as shown in Figure 1A. Only 2 channels are responsive in the insula, which are not shown here.

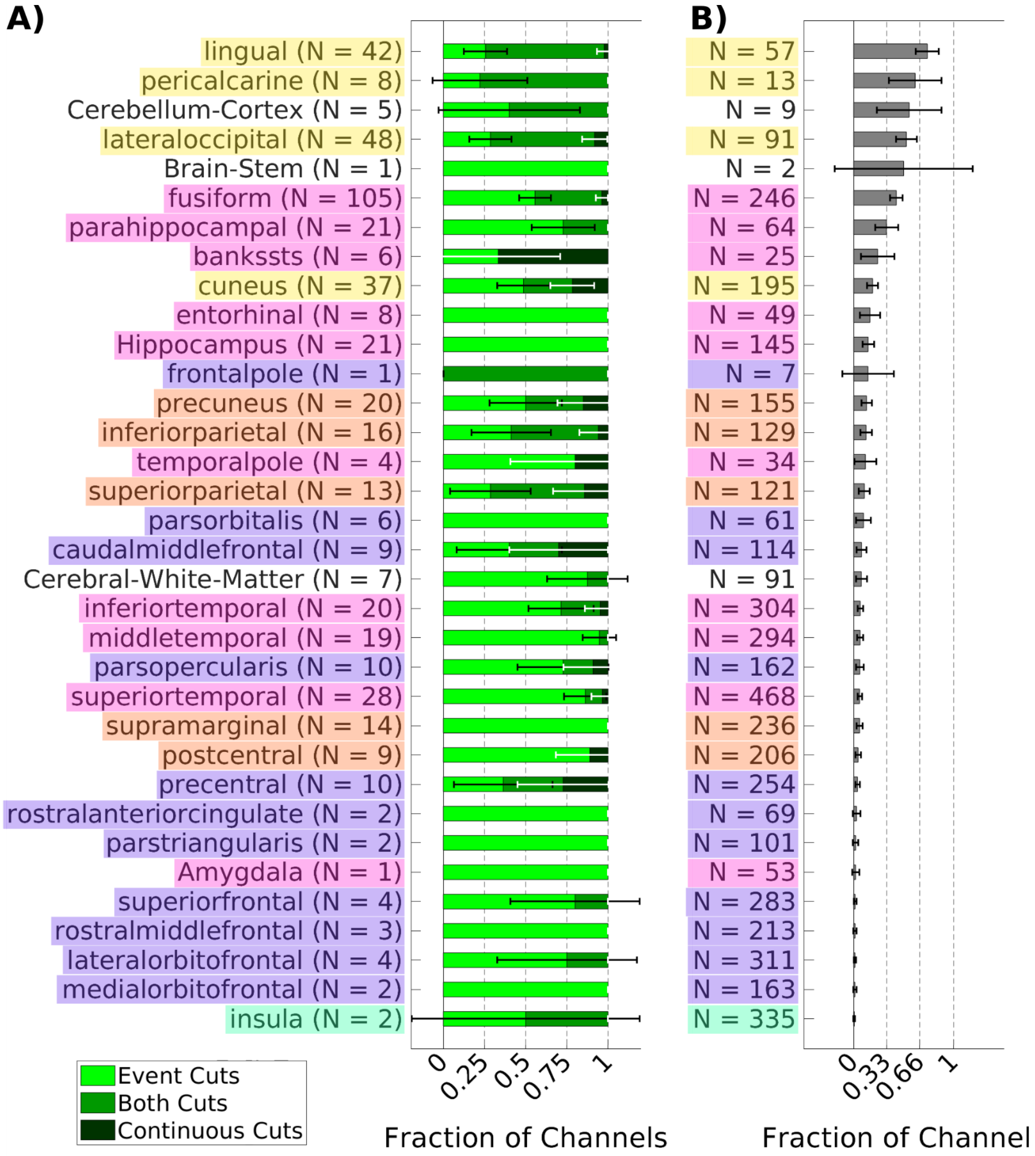

**Figure S12: Event cuts dominate neural responses in higher-order brain areas.** A) Fraction of channels responding to event cuts, continuous cuts, or both out of all responsive channels in regions of the Desikan-Killiany and Aseg atlas in freesurfer[3,4]. Error bars depict the 95% confidence interval of proportions. B) Fractions of channels responding to any time of film cuts out of all channels in each brain region.

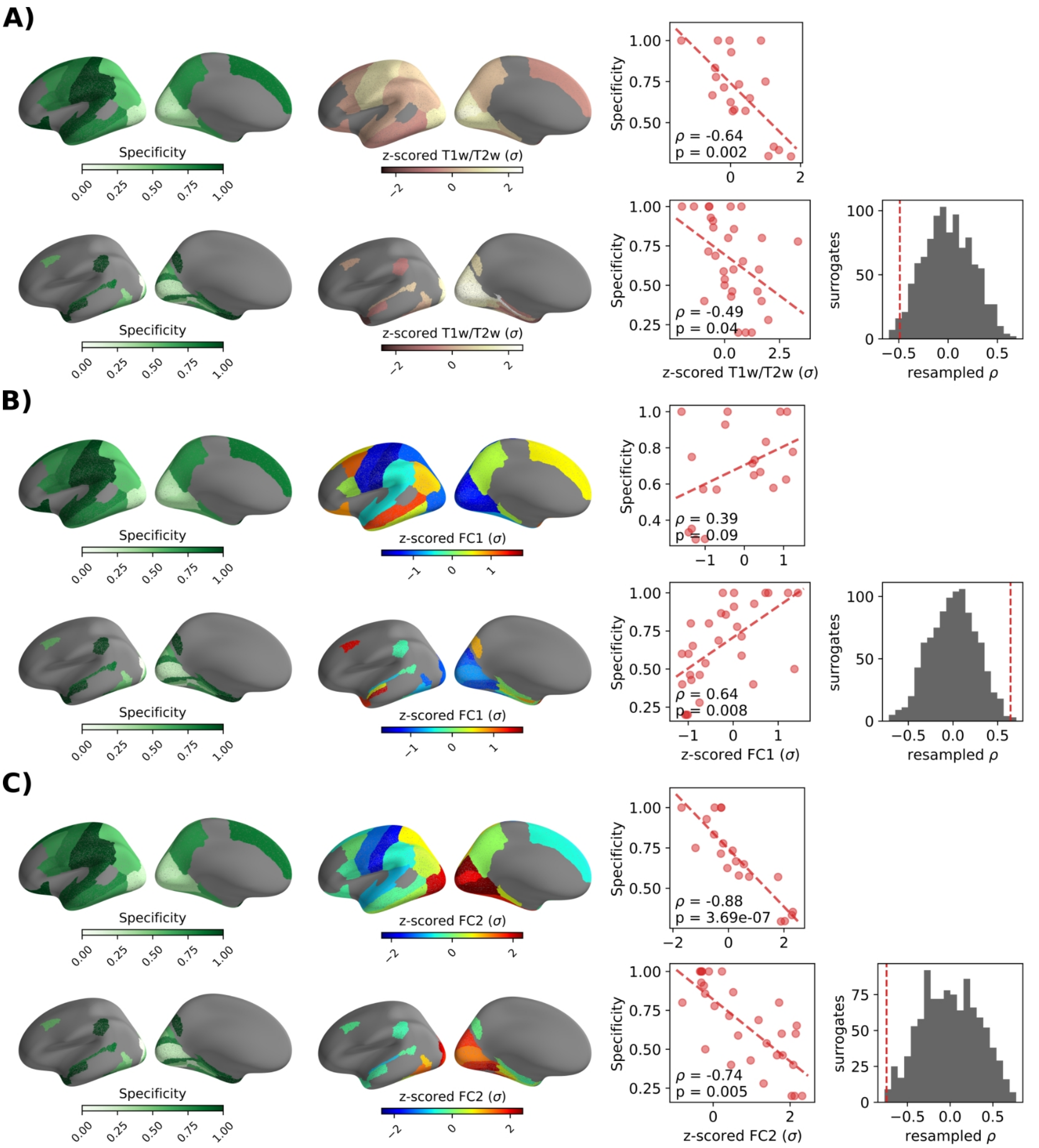

**Figure S13: Correlation of the specificity of responses to event versus continuous cuts with anatomical and functional gradients.** A) Top row: Spearman correlation between specificity of responses to event versus continuous cuts and T1w/T2w ratio in the Desikan-Killian atlas[3]. P-value is computed using the t-distribution. Bottom row: Spearman correlation between specificity and the T1w/T2w ratio in the Glasser atlas[9]. P-value is computed using permutation statistics by constructing a surrogate distribution of T1w/T2w ratio maps while preserving spatial autocorrelation following Gao et al.[10]. T1w/T2w ratio is a proxy of gray matter myelination that has been proposed to reflect a hierarchy from sensory to association areas[10,11]. B) Correlation of specificity with the first principal gradient of functional connectivity as in A. The first gradient of functional connectivity reflects a hierarchy from primary sensorimotor to transmodal regions[12]. C) Correlation of specificity to the second principal gradient of functional connectivity as in A. The second gradient of functional connectivity separates somatomotor and auditory cortices[12].

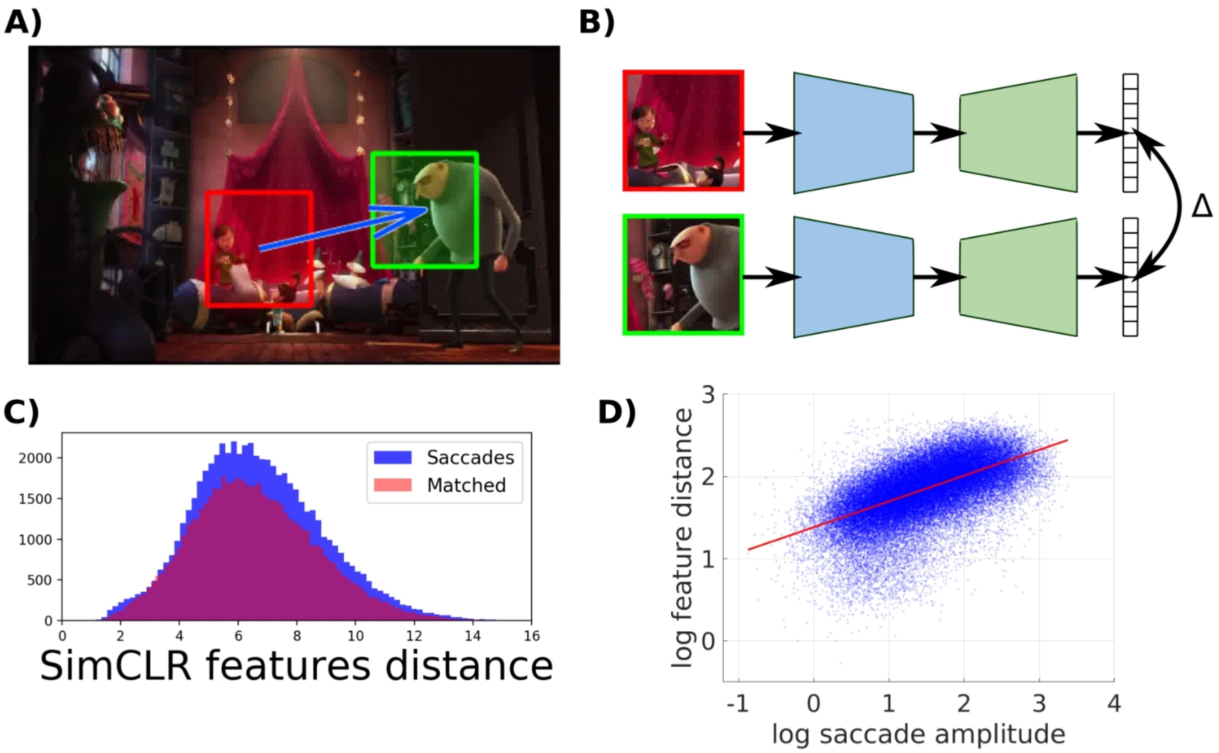

**Figure S14: Quantification of semantic novelty across saccades by contrastive learning.** A) Image patches with a size of 5 degree visual angle are cropped around the gaze point before (red box) and after (green box) saccades. B) Image patches are passed through a convolutional neural network pre-trained with contrastive learning to obtain a high-level representation in feature space. The distance Δ between these features quantifies the semantic novelty of saccades. C) Feature distance from image patches before and after saccades (blue), and distance between two patches from emulated saccades – random locations but matched to saccade amplitude and direction (red). Actual saccades have a larger distance than emulated saccades ($p=1.5*10^{-22}$, N=60170 saccades, N=60142 emulated saccades, unpaired Wilcoxon rank sum test). D) Correlation of feature distance with saccade amplitude. Saccades above and below the red line are selected as "high" and "low" novelty.

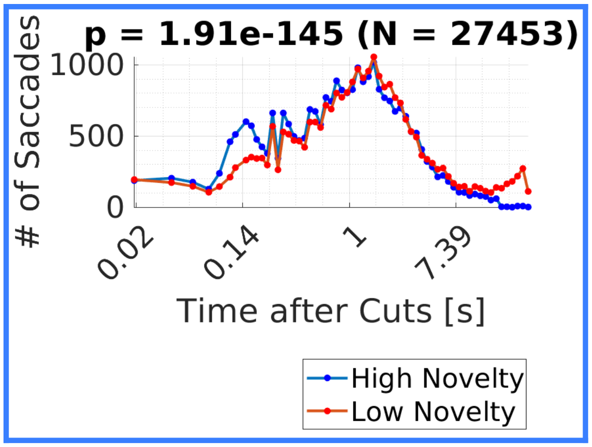

**Figure S15: Saccades with high novelty happen sooner after film cuts than saccades with low novelty.** The median of the time after the last film cut is 0.87s for saccades with high novelty and 1.12s for saccades with low novelty. The difference of 0.25s seconds between the median of both distributions is significant with p = $1.91*10^{-145}$ (N = 27453, Wilcoxon rank sum test). Note that the first saccade after film cuts has been removed from analysis to further mitigate interactions with cuts. Therefore, not many saccades are left in the first ~100 ms.

| **Factor** | **d.f.** | **F** | **p-value** |
|---|---|---|---|
| Condition | 1 | 19.77 | **$9.3*10^{-6}$** |
| Region | 5 | 20.41 | **$8.2*10^{-20}$** |
| Patient | 23 | 50.1 | **$1.6*10^{-170}$** |
| Error | 1632 | | |

**Table S6: Results of mixed-design ANOVA testing the significance of differences in response amplitudes to high and low novelty saccades (Figure 4A).** Condition is the fixed effect of amplitudes of high or low novelty saccades. Region is the fixed effect of brain region. Patient is a random effect.

| Factor | d.f. | F | p-value | | Factor | d.f. | F | p-value |
|---|---|---|---|---|---|---|---|---|
| **Occipital lobe** | | | | | **MTL** | | | |
| Condition | 1 | 9.06 | **0.0043** | | Condition | 1 | 0.43 | 0.56 |
| Patient | 12 | 14.72 | **$5.2*10^{-23}$** | | Patient | 15 | 13.68 | **$4.4*10^{-18}$** |
| Error | 264 | | | | Error | 105 | | |
| **Parietal lobe** | | | | | **Frontal lobe** | | | |
| Condition | 1 | 1.15 | 0.34 | | Condition | 1 | 3.37 | 0.09 |
| Patient | 18 | 18.31 | **$3.8*10^{-33}$** | | Patient | 21 | 27.96 | **$3.3*10^{-67}$** |
| Error | 210 | | | | Error | 445 | | |
| **Temporal lobe** | | | | | **Insula** | | | |
| Condition | 1 | 11.83 | **0.0011** | | Condition | 1 | 0.22 | 0.64 |
| Patient | 19 | 32.41 | **$3*10^{-74}$** | | Patient | 4 | 9.13 | **$2.5*10^{-4}$** |
| Error | 513 | | | | Error | 24 | | |

**Table S7: Results of mixed-design ANOVA testing the significance of differences in response amplitudes to high versus low novelty saccades within each brain region (Figure 4A).** Condition is the fixed effect of amplitudes of high or low novelty saccades. Patient is a random effect.

| | Occipital | Parietal | Temporal | MTL | Frontal | Insula |
|---|---|---|---|---|---|---|
| **Median Jaccard distance** | 0.2 | 0.41 | 0.69 | 0.67 | 1 | 1 |
| **p-value** | 0.999 | 0.999 | 0.6 | 0.999 | **0.024** | 0.55 |
| **N** | 23 | 14 | 18 | 3 | 15 | 1 |

**Table S8: Results of permutation statistics to test specificity of responses to high and low novelty saccades (Figure 4D).** 1,000 permutations. FDR corrected at q = 0.05. N: number of patients

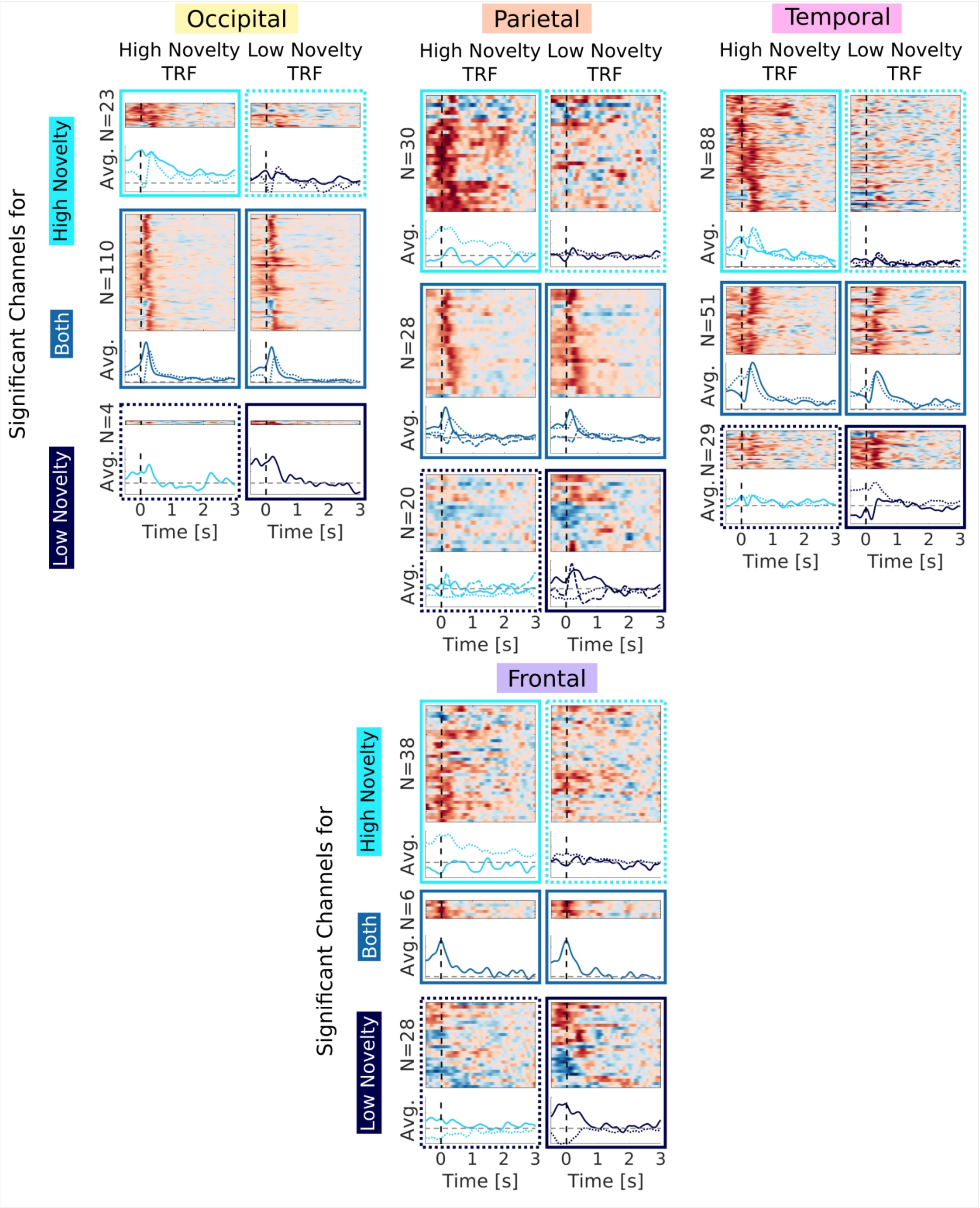

**Figure S16: Distinct temporal response functions to saccades with high or low novelty.** Separate filters are determined for saccades with high and low novelty (Figure 4C). In the top of each box normalized temporal response functions are depicted. The bottom of each box shows average temporal response functions. Averages are computed for clusters of similar temporal response functions. Light blue boxes show channels with significant responses to only high novelty saccades. Solid lines show the significant TRFs to high novelty saccades in these channels. Dashed lines show non-significant TRFs to low novelty saccades in the same channels. Dark blue boxes show TRFs in channels with significant responses to low novelty saccades. Solid lines, here, show the significant TRFs to low novelty saccades in these channels. Dashed lines show the non-significant TRFs to high novelty saccades in the same channels. Mixed blue boxes in the middle show channels with significant TRFs to both high and low novelty saccades. TRFs are grouped in lobes as shown in Figure 1A. Less than 20 channels are responsive in the MTL and insula, which are not shown here.

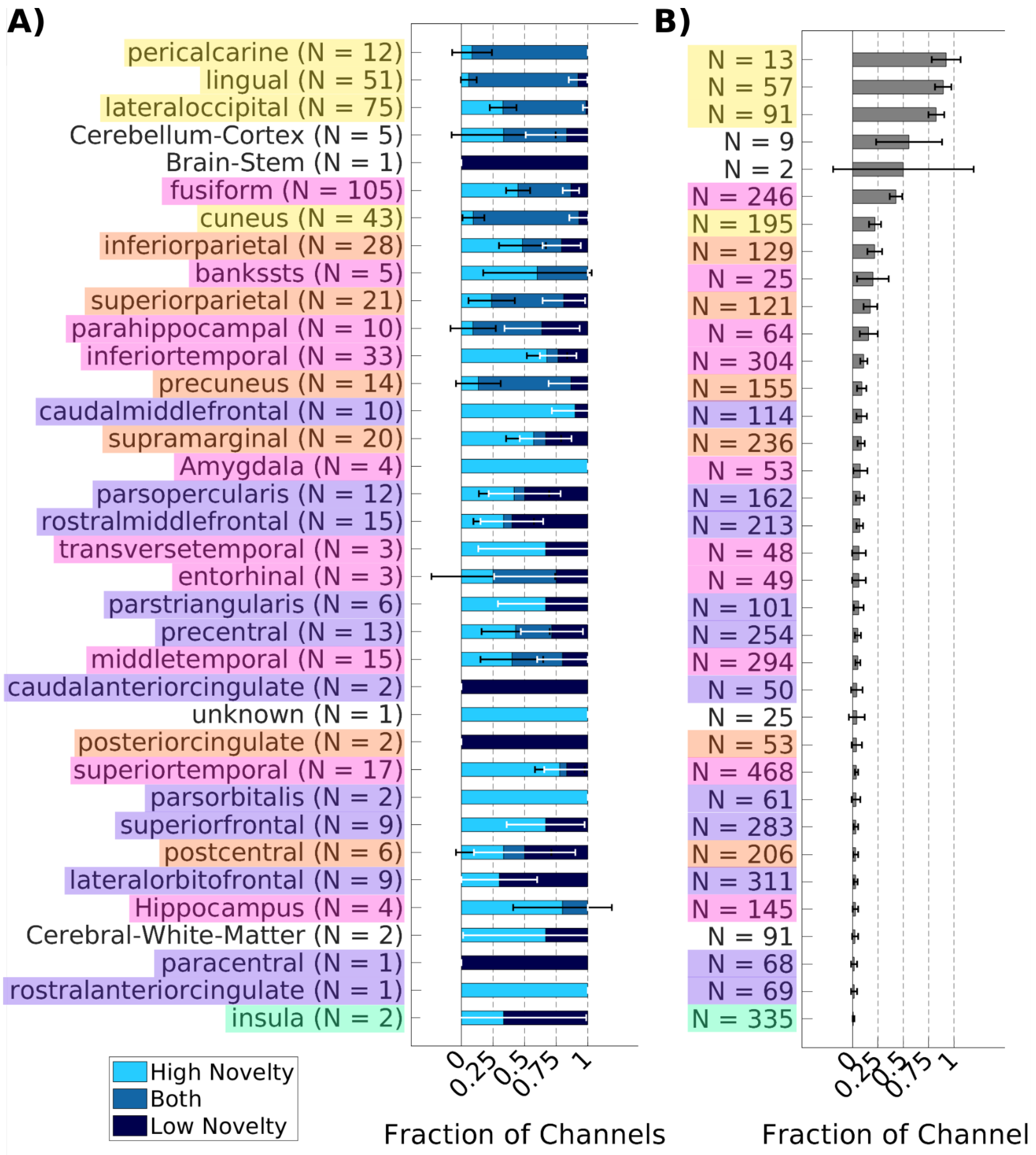

**Figure S17: Distinct sets of channels respond to saccades with high or low novelty.** A) Fraction of responsive channels out of all responsive channels. Channels are grouped in regions of the Desikan-Killiany and Aseg atlas in freesurfer[3,4]. In low-level visual areas such as the lingual gyrus, pericalcarine cortex and cuneus, most channels have significant TRFs to saccades with both high and low novelty. Higher order brain areas show more specific responses to either type of saccade. Notably, the majority of channels in the amygdala respond only to saccades with high novelty. Channels in areas of the frontal lobe, such as the lateral orbitofrontal and rostral middle frontal cortex respond to either saccades with high or low novelty only. B) Fraction of channels with significant responses to either saccades with high or low novelty out of all channels in each brain area.

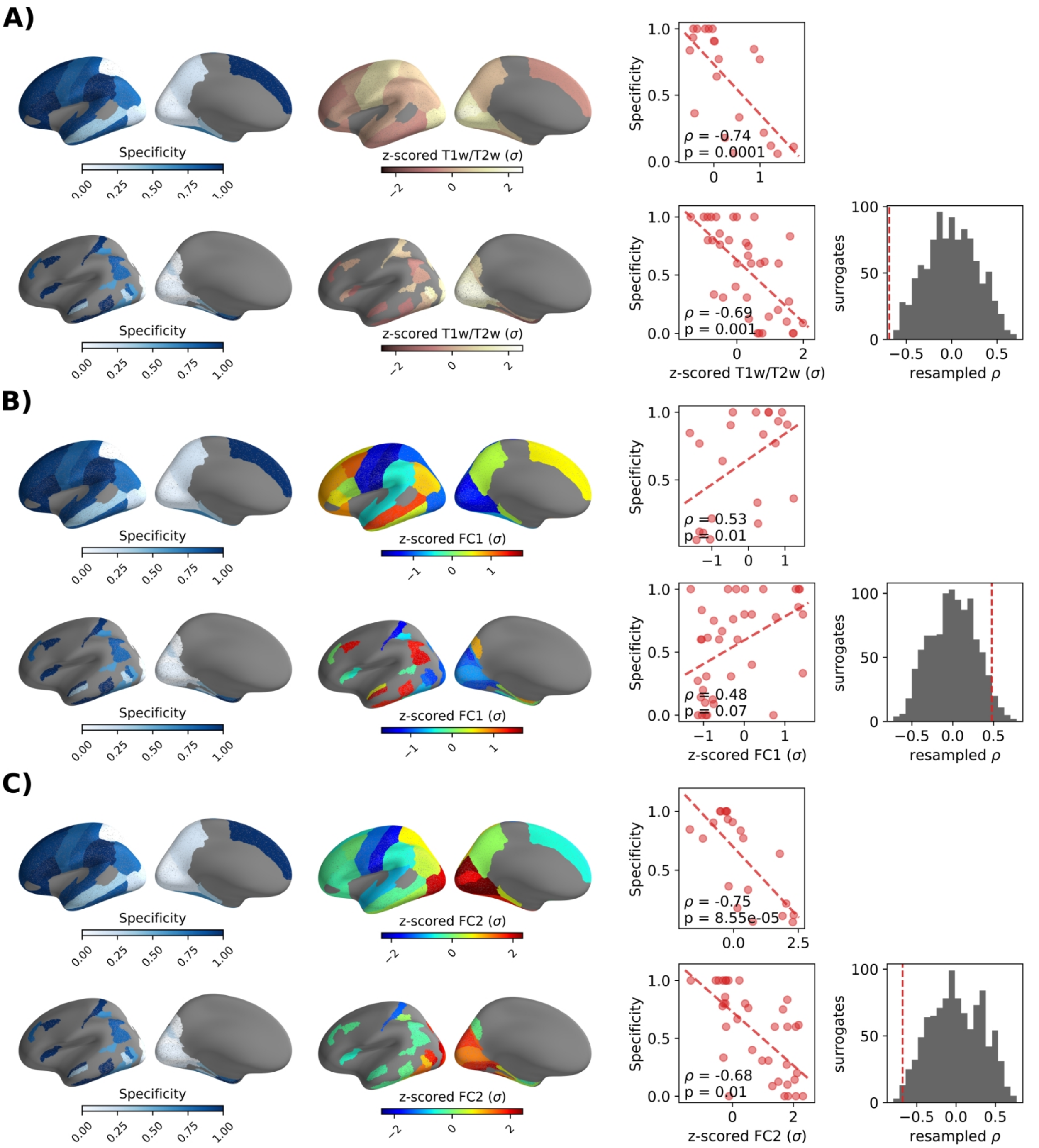

**Figure S18: Correlation of the specificity of responses to high- versus low-novelty saccades with anatomical and functional gradients.** A) Top row: Spearman correlation between specificity of responses to high- versus low-novelty saccades and T1w/T2w ratio in the Desikan-Killian atlas[3]. P-value is computed using the t-distribution. Bottom row: Spearman correlation between specificity and the T1w/T2w ratio in the Glasser atlas[9]. P-value is computed using permutation statistics as describes in Figure S13. B) Correlation of specificity with the first principal gradient of functional connectivity as in A. C) Correlation of specificity to the second principal gradient of functional connectivity as in A.

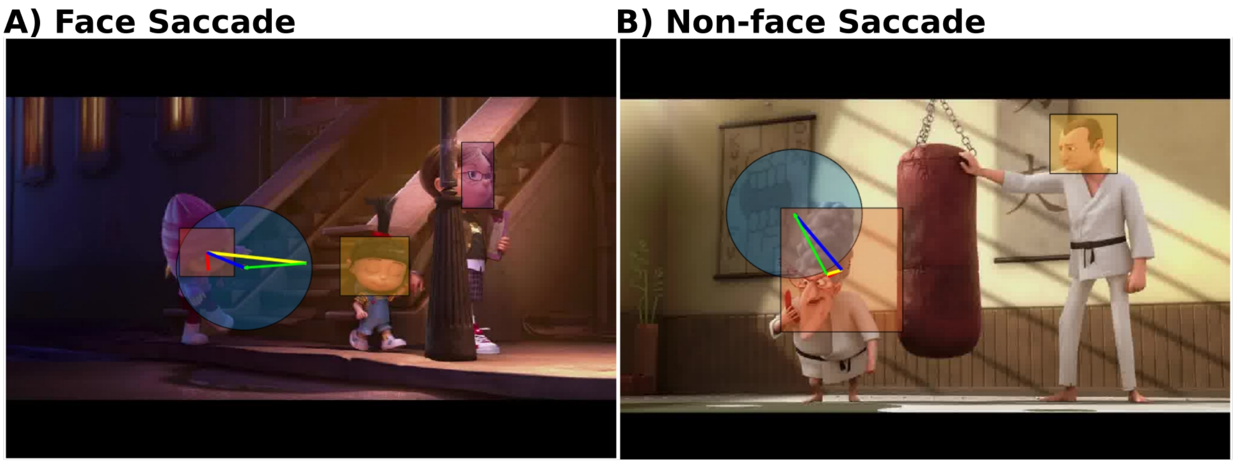

**Figure S19: Classification of saccades in face and non-face saccades.** Green vector: saccade; red circle: fixation onset; yellow line: vector from saccade onset to centroid of the closest face annotation bounding box; blue line: vector from fixation onset to centroid of the closest face annotation bounding box; red line: normal vector to the saccade vector intersecting with the face annotation centroid; blue circle: area of the foveal image of 5 degree visual angle around the fixation onset; yellow, orange and purple boxes: face annotation bounding boxes. Features for face detection: 1. Saccade direction: 1 towards face, -1 away from face. Direction of the green vector in relation to the closest bounding box. 2. Face area: Intersection between blue circle and bounding box in mm2. 3: Distance to face: Distance of blue line from fixation onset to bounding box centroid in mm. 4. Saccade angle: Angle between green saccade vector and yellow line. 5. Centroid angle: Angle between yellow and blue vector. A) Face saccade with fixation onset just outside the face annotation bounding box. Features: 1. Saccade direction: 1; 2. Face area: 552$mm^2$; 3. Distance to face: 69mm; 4. Saccade angle: 11°; 5. Centroid angle: 17°. B) Non-face saccade with fixation onset within a face annotation bounding box. Features: 1. Saccade direction: -1; 2. Face area: 994$mm^2$; 3. Distance to face: 127mm; 4. Saccade angle: 99°; 5. Centroid angle: 69°

| **Factor** | **d.f.** | **F** | **p-value** |
|---|---|---|---|
| Condition | 1 | 40.4 | **$2.7*10^{-10}$** |
| Region | 5 | 11.46 | **$7*10^{-11}$** |
| Patient | 22 | 25.99 | **$9.7*10^{-90}$** |
| Error | 1551 | | |

**Table S9: Results of mixed-design ANOVA testing the significance of differences in response amplitudes to face and non-face saccades (Figure 5A).** Condition is the fixed effect of amplitudes of face or non-face saccades. Region is the fixed effect of brain region. Patient is a random effect.

| Factor | d.f. | F | p-value | | Factor | d.f. | F | p-value |
|---|---|---|---|---|---|---|---|---|
| **Occipital lobe** | | | | | **MTL** | | | |
| Condition | 1 | 30.84 | **$1.6*10^{-7}$** | | Condition | 1 | 9.26 | **0.0036** |
| Patient | 11 | 5.18 | **$4.5*10^{-7}$** | | Patient | 14 | 7.88 | **$1.6*10^{-10}$** |
| Error | 263 | | | | Error | 100 | | |
| **Parietal lobe** | | | | | **Frontal lobe** | | | |
| Condition | 1 | 19.01 | **$3.6*10^{-5}$** | | Condition | 1 | 13.43 | **$3.7*10^{-4}$** |
| Patient | 16 | 10.09 | **$8*10^{-18}$** | | Patient | 20 | 18.13 | **$2.6*10^{-44}$** |
| Error | 200 | | | | Error | 432 | | |
| **Temporal lobe** | | | | | **Insula** | | | |
| Condition | 1 | 0.08 | 0.84 | | Condition | 1 | 0.04 | 0.85 |
| Patient | 18 | 14.00 | **$7.2*10^{-33}$** | | Patient | 5 | 9.76 | **$4.3*10^{-5}$** |
| Error | 464 | | | | Error | 25 | | |

**Table S10: Results of mixed-design ANOVA testing the significance of differences in response amplitudes to face versus non-face saccades within each brain region (Figure 5A).** Condition is the fixed effect of amplitudes of face or non-face saccades. Patient is a random effect.

| | Occipital | Parietal | Temporal | MTL | Frontal | Insula |
|---|---|---|---|---|---|---|
| **Median Jaccard distance** | 0.46 | 0.94 | 0.96 | 0.4 | 1 | 1 |
| **p-value** | 0.999 | **0.002** | **0.002** | 0.999 | **0.002** | **0.023** |
| **N** | 12 | 12 | 17 | 5 | 15 | 2 |

**Table S11: Results of permutation statistics to test specificity of responses to face and non-face saccades (Figure 5D).** 1,000 permutations. FDR corrected at q = 0.05. N: number of patients

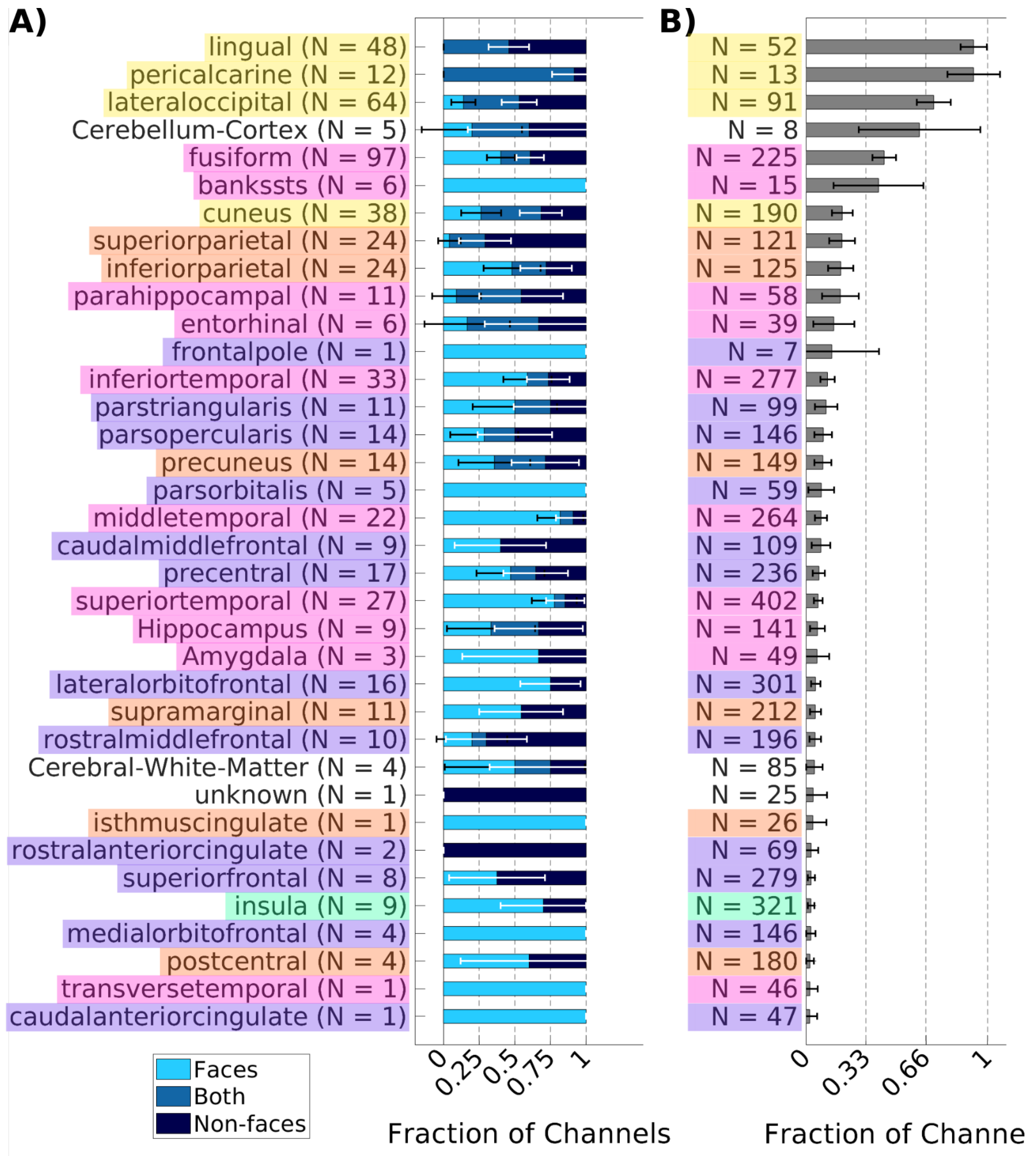

**Figure S20: Distinct sets of channels respond to face and non-face saccades.** A) Fraction of channels responding to face or non-face saccades out of all responsive channels in each area of the Desikan-Killiany and Aseg atlas in freesurfer[3,4]. Channels in the transverse temporal gyrus (Heschl's gyrus) and superior temporal gyrus of the auditory cortex respond almost exclusively to face saccades only. B) Fraction of channels responding to either face or non-face saccades out of all channels in each brain area.

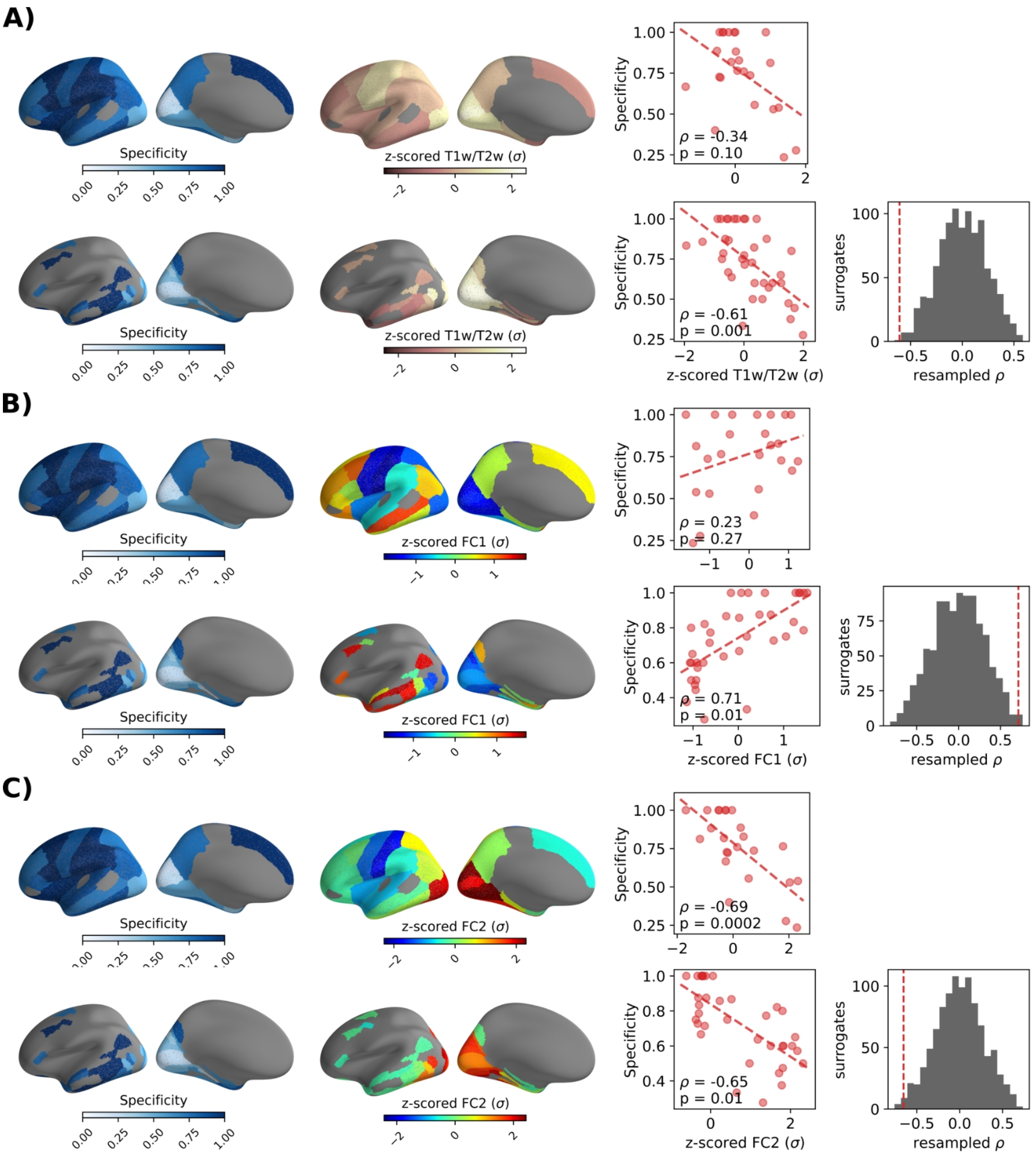

**Figure S21: Correlation of the specificity of responses to saccades to face versus non-face objects with anatomical and functional gradients.** A) Top row: Spearman correlation between specificity of responses to face versus non-face saccades and T1w/T2w ratio in the Desikan-Killiany atlas[3]. P-value is computed using the t-distribution. Bottom row: Spearman correlation between specificity and the T1w/T2w ratio in the Glasser atlas[9]. P-value is computed using permutation statistics as describes in Figure S13. B) Correlation of specificity with the first principal gradient of functional connectivity as in A. C) Correlation of specificity to the second principal gradient of functional connectivity as in A.

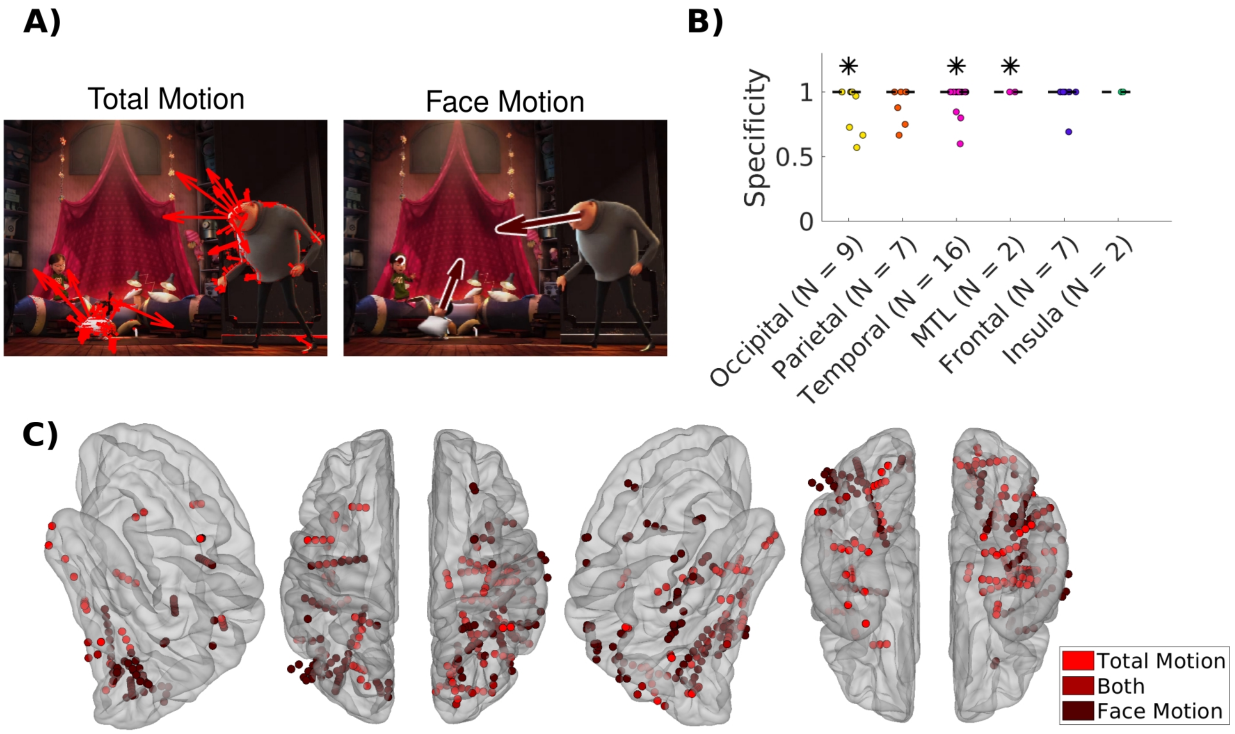

**Figure S22: Total motion and face motion are processed in distinct visual areas.** A) Separate TRFs are computed for total motion (optical flow) and face motion, capturing socially relevant information. Regressors for film cuts and saccades are included to control for correlated activity. To estimate face motion we compute the velocity of the centroid of the face annotations from frame to frame. We sum the velocity of all bounding boxes within each frame to capture motion of all faces within a frame. B) Specificity of responses to total motion and face motion as defined as in Figure 3D. C) Channels with significant response to total motion only (bright red), face motion only (dark red), and both total and face motion (medium red) on the fsaverage brain. For results in a more detailed parcellation of the brain see Figure S23. Channel locations are plotted on the freesurfer fsaverage brain[5] with the iELVis MATLAB toolbox[6].

| | **Occipital** | **Parietal** | **Temporal** | **MTL** | **Frontal** | **Insula** |
|---|---|---|---|---|---|---|
| **Median Jaccard distance** | 1 | 1 | 1 | 1 | 1 | 1 |
| **p-value** | **0.036** | 0.12 | **0.006** | **0.036** | 0.2 | 0.14 |
| **N** | 9 | 7 | 76 | 2 | 7 | 2 |

**Table S12: Results of permutation statistics to test specificity of responses to face motion and total motion (Figure S22B).** 1,000 permutations. FDR corrected at q = 0.05. N: number of patients

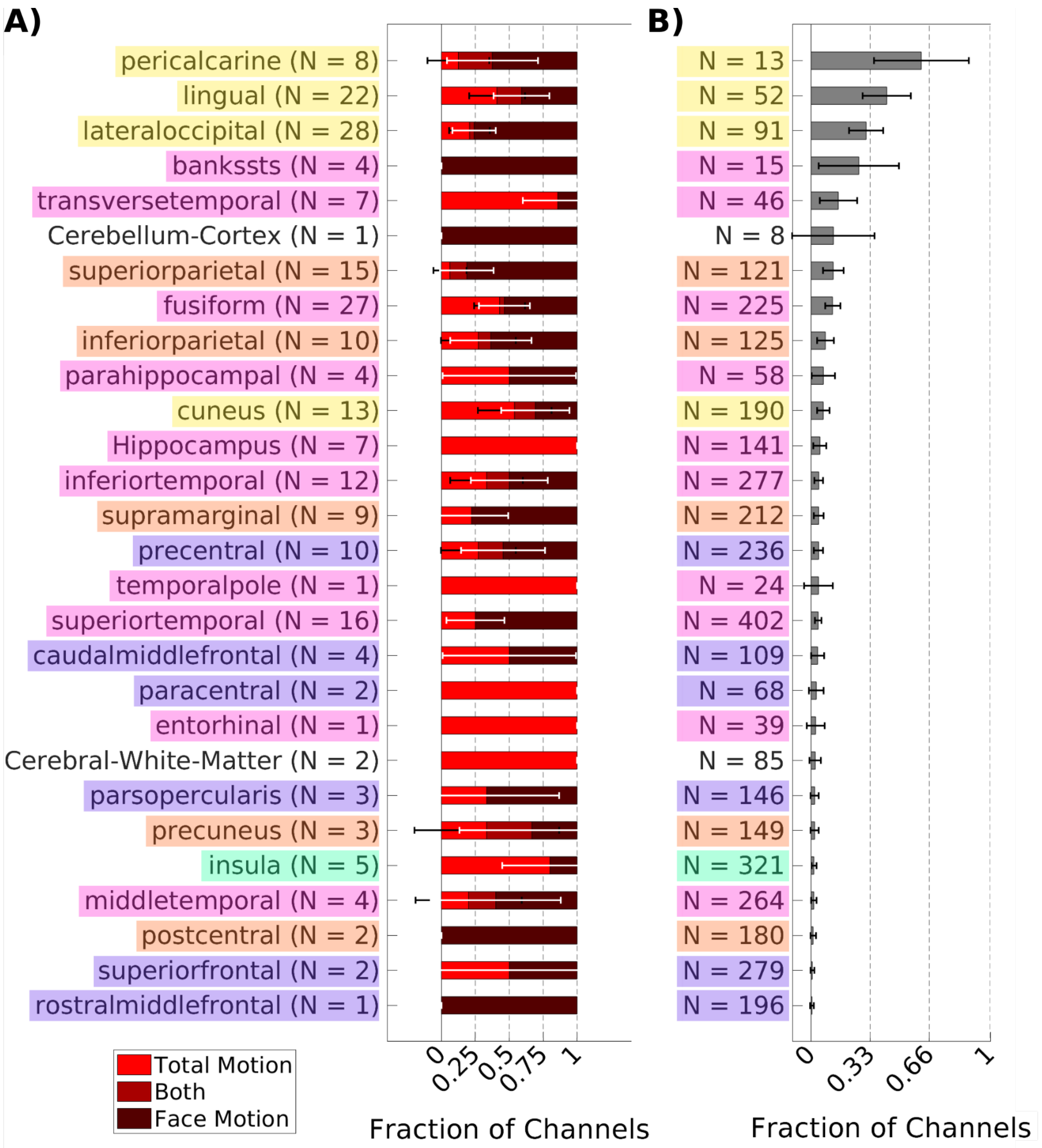

**Figure S23: Responses to total motion are more widespread than responses to face motion.** A) Fraction of channels responding to total motion (optical flow) or face motion out of all responsive channels in each area of the Desikan-Killiany and Aseg atlas in freesurfer[3,4]. In most brain areas more channels respond to total motion than face motion. Exceptions are areas that are known to be involved in processing faces, such as the lateral occipital cortex and fusiform gyrus. In these areas more electrodes respond to face motion than total motion. B) Fraction of channels responding to either total motion or face motion out of all channels in each brain area.

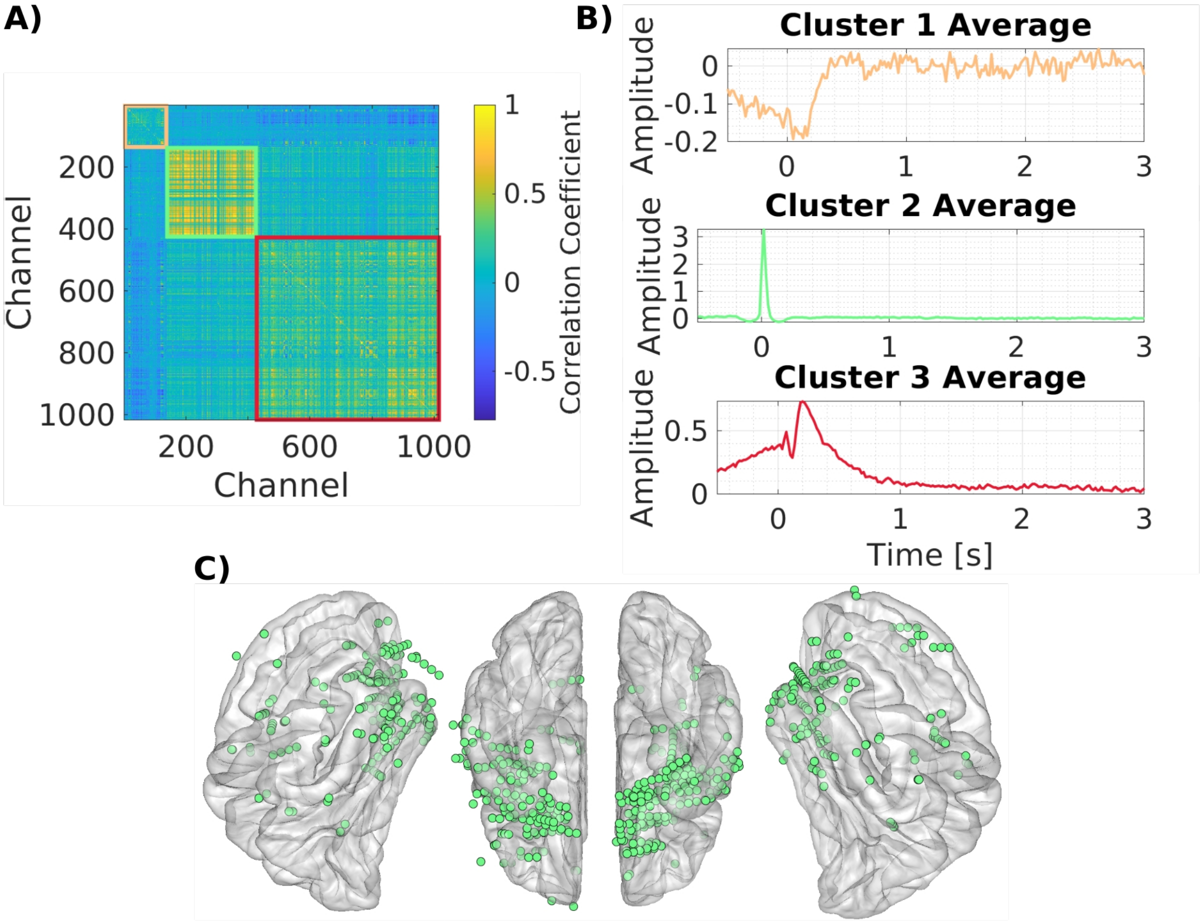

**Figure S24: Detection and removal of saccadic artifacts.** A) Correlation matrix of all significant TRFs to saccades (Figure 2 & S4A). The correlation matrix was grouped in clusters of channels with high correlation to each other[7]. Three clusters have been chosen a-priori. Visually the data is well organized with 3 clusters. B) Average TRFs for all clusters. Cluster 1 contains the saccade artifact, which is visually identified. All 466 channels belonging to Cluster 1 are removed from the analysis (Figure 2 & S3A). C) Location of all channels belonging to Cluster 1 on the fsaverage brain. Most channels are located around the orbit of the eye and thus likely contain muscular artifacts related to saccade execution. Channel locations are plotted on the freesurfer fsaverage brain[4,5] with the iELVis MATLAB toolbox[6].
